# Activation of the muscle-to-brain axis ameliorates neurocognitive deficits in an Alzheimer’s disease mouse model via enhancing neurotrophic and synaptic signaling

**DOI:** 10.1101/2024.06.14.599115

**Authors:** Hash Brown Taha, Allison Birnbaum, Ian Matthews, Karel Aceituno, Jocelyne Leon, Max Thorwald, Jose Godoy-Lugo, Constanza J. Cortes

## Abstract

**INTRODUCTION:** Skeletal muscle regulates central nervous system (CNS) function and health, activating the muscle-to-brain axis through the secretion of skeletal muscle originating factors (‘myokines’) with neuroprotective properties. However, the precise mechanisms underlying these benefits in the context of Alzheimer’s disease (AD) remain poorly understood.

**METHODS:** To investigate muscle-to-brain axis signaling in response to amyloid β (Aβ)- induced toxicity, we generated 5xFAD transgenic female mice with enhanced skeletal muscle function (5xFAD;cTFEB;HSACre) at prodromal (4-months old) and late (8-months old) symptomatic stages.

**RESULTS:** Skeletal muscle TFEB overexpression reduced Aβ plaque accumulation in the cortex and hippocampus at both ages and rescued behavioral neurocognitive deficits in 8- months-old 5xFAD mice. These changes were associated with transcriptional and protein remodeling of neurotrophic signaling and synaptic integrity, partially due to the CNS-targeting myokine prosaposin (PSAP).

**DISCUSSION:** Our findings implicate the muscle-to-brain axis as a novel neuroprotective pathway against amyloid pathogenesis in AD.

## 1. BACKGROUND

Multiple studies have demonstrated that the loss of neurotrophic signaling and synaptic connections are early events during Alzheimer’s disease (AD) pathogenesis. Indeed, deficits in neurotrophic factor abundance alongside widespread changes in synapse number, structure and function have been reported in the CNS of persons with, and animal models of, AD [1–4]. Furthermore, these deficits correlate well with cognitive abnormalities in persons with AD, even more so than other pathological markers of the disease [1, 3]. This suggests that targeting neurotrophic signaling or the synapse may be a critical step in developing new generations of AD disease-modifying therapeutics.

The muscle-to-brain axis has recently arisen as a powerful regulator of CNS health and function [5–9]. This novel signaling axis is mediated by circulating factors collectively known as ‘myokines’, which are secreted from skeletal muscle in response to multiple physiological and pathological conditions with high metabolic demand [10]. Through mechanisms that remain poorly understood, these circulating myokines can promote CNS rejuvenation and neuroprotection, reducing neuroinflammation, reactivating declining neurogenesis and reducing the accumulation of protein aggregates [5, 11–15], all key features of AD pathogenesis. However, to date, the precise mechanisms that facilitate myokine-dependent neuroprotection in the context of AD remain poorly understood.

We have previously shown that activation of the muscle-to-brain axis via targeted overexpression Transcription Factor E-B (TFEB) in skeletal muscle can protect against age-associated cognitive decline and reduce degenerative neuropathological hallmarks in the MAPT P301S mouse model of tauopathy [5]. Indeed, we observed robust reductions in neuroinflammation and rescue of declining neurotrophic signaling in the hippocampus of both aged (24-months-old) and MAPT P301S transgenic mice (8-months-old) with muscle-TFEB overexpression. Furthermore, we also identified sex-dimorphic transcriptional remodeling in the CNS of muscle-TFEB overexpressing mice, with female mice displaying significant up-regulation in pathways associated with synaptic physiology, neurotransmitter signaling and voltage- and ion-gated channels, and male mice displaying up-regulation of mitochondrial and ribosomal signaling [5]. We also confirmed that these neuroprotective effects were associated with increased expression and secretion of known CNS-targeting myokine cathepsin B [5]. However, whether muscle-TFEB overexpression in skeletal muscle impacts CNS health in the presence of Aβ degenerative toxic species has yet to be explored.

Here, we report that activation of the muscle-to-brain axis using our cTFEB;HSACre (muscle-TFEB) mice also provides important neuroprotective effects in the 5xFAD model of Aβ-plaque toxicity. Muscle-TFEB overexpression reduced Aβ plaque accumulation in the cortex and hippocampus at prodromal (4-months-old) and later stages (8-months-old) of disease. Importantly, muscle-TFEB rescued neurocognitive deficits in 5xFAD mice in a battery of memory and learning behavioral tests. These neuroprotective effects appear to be independent of modulation of neuroinflammatory states. Instead, muscle-TFEB expression promoted transcriptional remodeling of the 5xFAD CNS in pathways associated with neurotrophic and synaptic signaling, which we validate via extensive immunoblotting for neurotrophic and synaptic markers. Furthermore, we also provide evidence demonstrating our newly identified exercise-responsive myokine prosaposin (PSAP) as a potential mechanism driving these benefits.

Our results indicate that the promotion of an exercise-like state in skeletal muscle via TFEB-overexpression is sufficient to provide neuroprotection against amyloid pathogenesis and neurocognitive decline by enhancing neurotrophic and synaptic signaling, potentially through the release of CNS-targeting myokines such as PSAP. These findings support the neuroprotective effects of the muscle-to-brain axis and provide potential pathways that can be targeted by interventions to preserve cognitive function and prevent worsening pathologies against neurodegenerative diseases.

## 2. METHODS

### 2.1 Animals

We have previously described the generation of fxSTOP-TFEB transgenic mice [5]. For this work, cTFEB;HSACre;5xFAD transgenic mice were generated by crossing fxSTOP-TFEB females with 5xFAD (MMRRC Strain: 034840-JAX | B6SJL Tg(APPSwFlLon,PSEN1*M146L*L286V)6799Vas/Mmjax) males. Double transgenic females were then crossed with heterozygous HSACre males (JAX Strain No: 006139 | HSACre79) (**Figure 1A**). All lines have been maintained in a C57B6/L background for over ten generations in our lab. Mice were housed in a temperature- and humidity-controlled environment on a 12- hour light/dark cycle with food and water ad libitum. For all experiments, control animals were TFEB-;HSACre+ and/or TFEB+;HSACre-littermates, which we have previously demonstrated have no neurological or metabolic phenotypes at this age [5]. To our knowledge, we did not detect any differences in plaque-associated CNS phenotypes in 5xFAD+;TFEB+ or 5xFAD+;HSACre+ double transgenic animals compared to 5xFAD single transgenic controls (**Figures 1-5 and S1-5**), and are thus all grouped into the ‘AD’ 5xFAD group. All animal experimentation adhered to NIH guidelines and was approved by and performed in accordance with the University of Southern California and University of Alabama at Birmingham Institutional Animal Care and Use.

**FIGURE 1.**
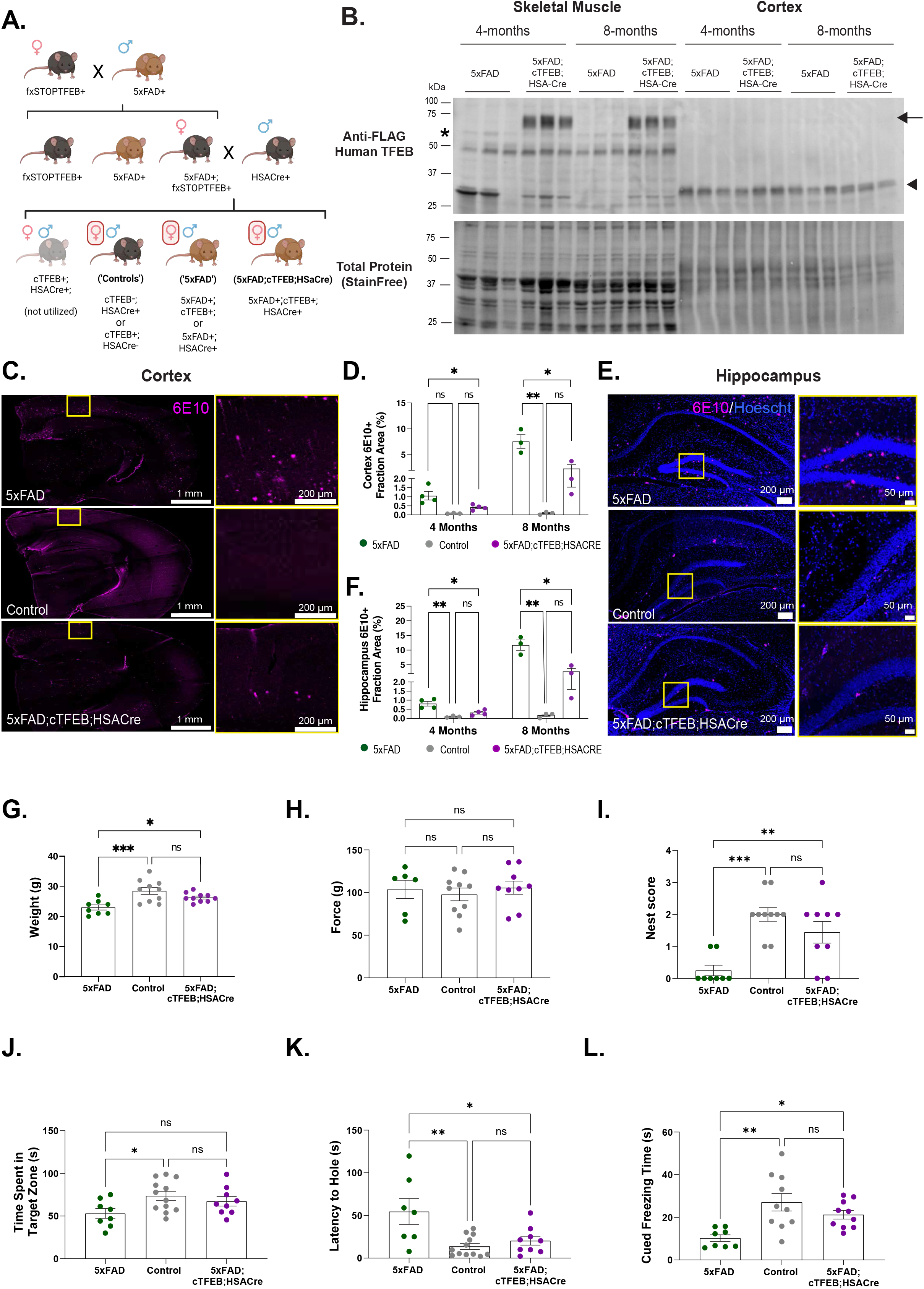
Skeletal muscle-TFEB overexpression reduces amyloid β (Aβ) accumulation and rescues behavioral neurocognitive deficits in 5xFAD 4- and 8-months old transgenic female mice. **(A)** Schematic depicting the breeding scheme to generate 5xFAD;cTFEB;HSACre animals. The bottom row depicts animals used in the study. Control animals are negative for 5xFAD (cTFEB-;HSACre+ or cTFEB+;HSACre-), AD animals are positive for 5xFAD (5xFAD+;cTFEB+;HSACre- or 5xFAD+;cTFEB-;HSACre+), and 5xFAD;cTFEB;HSACre mice are positive for all 3 transgenes. **(B)** Immunoblots of skeletal muscle and brain from 5xFAD and 5xFAD;cTFEB;HSACre 4- and 8-months-old mice confirming 3x-FLAG-TFEB protein expression in skeletal muscle but not brain of 5xFAD;cTFEB;HSACre. Transgenic 3x-FLAG-TFEB protein weighs ∼75 kDa. Arrow highlights TFEB protein, asterisk (*) highlights a non-specific band, arrowhead highlights the FLAG-eGFP fragment present in TFEB+ individuals [5]. Total protein stain-free lanes are shown below as a loading control. Densitometry quantification relative to Total StainFree protein lane densitometry is shown in **Figure S1A**. **(C)** Representative images of coronal sections from 4-months-old mice stained for Aβ plaques (6E10, magenta). **(D)** Quantification of cortical Aβ plaque load for 4- and 8-months-old mice. **(E)** Representative images of the hippocampus from coronal sections of 4-months-old mice stained for Aβ plaques (6E10, magenta). **(F)** Quantification of hippocampal Aβ plaque load for 4- and 8-months-old mice. Each point represents the average of all anterior (n = 3), medial (n = 5) and posterior (n = 3) sections containing the cortex or hippocampus for an individual animal. Scale bars as shown. **(G-L)** Behavioral and neurocognitive phenotyping of 8-months-old mouse cohorts **(G)** Body weight, **(H)** Grip strength, **(I)** Nest building scores, **(J-K)** Barnes maze testing evaluating the time spent in the target zone **(J)** and latency to the target hole **(K)**. **(L)** Fear conditioning evaluating freezing time during cue-dependent trials. Statistical comparison was performed using one-way ANOVA and post hoc multiple comparisons, ^∗^p < 0.05, ^∗∗^p < 0.01, ^∗∗∗^p < 0.001, n.s. non-significant. Data is represented as mean ± SEM.

### 2.2 Tissue Collections

Animals were anesthetized with 3.8% Avertin Solution and received transcardial perfusion with 60 mL of chilled 1X PBS. Right hemibrains were post-fixed in 4% paraformaldehyde for less than 24 hours before transferring to a 30% (w/v) sucrose solution for another 24 hours before OCT (TissueTek, 4583) embedding and cryo-sectioning. The left hemisphere was micro-dissected into cortex and hippocampus and flash-frozen in liquid nitrogen alongside quadricep skeletal muscle for downstream analysis. Blood was collected via cardiac puncture and transferred to an EDTA tube in a 1:100 ratio and centrifuged at 1,500xg for 15 minutes at 4°C to separate the layers. The top layer containing plasma was transferred to a 1.5 mL tube. The other layers were discarded. Tissue and plasma samples were stored at -80°C prior to downstream analysis.

### 2.3 Immunofluorescence Staining and Analysis

Tissue was embedded in OCT (TissueTek, 4583), slow frozen using 2-methylbutane vapors (VWR, 103525-278) submerged in liquid nitrogen and stored at -80°C until use. All hemibrains were sectioned on a standing cryostat in 20 μm slices. All sections were permeabilized with 1X PBS + 0.1% (v/v) Triton for 15 minutes and blocked with 4% (w/v) BSA in 1X PBS for 1 hour. Sections were stained with appropriate antibodies overnight at 4°C. See **Table S1** for full details on antibodies utilized in this project. After overnight incubation, sections were washed three times with 1X PBS and incubated with appropriate secondary antibodies for 1 hour. Sections were washed three times with 1X PBS and incubated with 1:5000 Hoescht (Thermo Scientific, 62249) for 5 minutes. Lastly, sections were washed three times with 1X PBS and mounted with prolong glass (Invitrogen, P36984). Slides were imaged using a 20X or 60X objective on the Nikon A15R/SIM Confocal microscope at the UAB HRIF Confocal/Light Microscopy Core, or on Stellaris 5 (Leica Microsystems, 8119637).

Aβ plaque load was quantified following an established protocol [16] in ImageJ. Images included at least three biological replicates from each group and at least eleven technical replicates from anterior (n = 3), middle (n = 5) or posterior (n = 3) coronal sections. Cortical quantification spanned all six layers of the cortex and the whole morphometric structure of the hippocampus. Relative presence of glial cells in brain cryosections of the dentate gyrus of the hippocampus was quantified by the percentage of area coverage of GFAP+ (astrocyte) and IBA1+ (microglia) staining using ImageJ. The number of astrocytes and the volume of microglia as a proxy for neuroinflammation was determined using the Surfaces function available through Imaris Microscopy Image Analysis Software. Images included at least 3 biological replicates from the 5xFAD and 5xFAD;cTFEB;HSACre groups, and 2-3 biological replicates from controls. All measurements included at least three technical replicates from middle coronal sections.

### 2.3 Mouse Phenotyping and Behavior

#### 2.3A Open Field

Mice (8-months-old) were individually placed in an open field (50 x 50 cm) illuminated to 600 lux in the center, and were allowed to explore freely for 5 minutes. The following behaviors were recorded: traveled distance (cm), time spent in the center (s) and velocity (cm/s). Movements were tracked via Ethovision (Noldus).

#### 2.3B Nest Building

Nest building was assessed following similarly established protocols [17]. In brief, mice were transferred to individual testing cages with wood-chip bedding but no environmental enrichment items such as paper towel or other nesting material, and one (∼3g) piece of nestlet was placed in each cage. The nests were assessed at 24 hours after on a rating scale of 0 (no nestlet interaction) to 5 (fully developed nest) based on nest construction.

#### 2.3C Barnes Maze

The maze was an open circular platform (75 cm in diameter), with 20 evenly spaced holes located 5 cm apart along the border with a rectangular escape box (5.5 × 11.5 × 5.5 cm) placed under one of the holes. Distinct and constant spatial cues were located around the maze. Mice completed each trial of the maze individually. On the first day, a red light was turned on, the mice were placed in the center inside a start box for 10 seconds, and were allowed to explore the maze for 3 minutes. On the second day, an initial training occurred where mice were placed inside the start box for 10 seconds with the buzzer and light on (80 dB + 400 lux) and were guided to the escape box, where the mice remained for an additional minute before their return to the home cage. Following this, the first training session began by placing the mice inside the start box for 10 seconds with the buzzer and light on. The mice were allowed to explore for 3 minutes or until the mice found the escape box; once they found the escape, they remained in the box for an additional minute. Mice were returned to the home cage after testing ends. Days 3-5 were the same as Day 2, but did not include the initial training guide. On Day 7, the escape box was converted into a decoy box; the mice were placed in the center of maze inside the start box for 10 seconds with the buzzer and light on, and were allowed to explore for 3 minutes. Video recordings and data were analyzed with Ethovision (Noldus).

#### 2.3D Fear Conditioning

Conditioning was carried out in Plexiglas Freeze Monitor chambers (26 cm × 26 cm × 17 cm) with speakers and lights mounted on two opposite walls, and shockable grid floors. The chambers were housed in sound proofed boxes. Sessions were recorded with real-time digital video and recordings were calibrated to distinguish among subtle movements, such as whisker twitches, tail flicks, and freezing behavior. On Day 1, mice were habituated to the chambers in a 5 min shock-free test. On Day 2, the mice are subjected to the context and conditioned stimulus (30 seconds, 2800 Hz, 75 dB sound + white light) in association with foot shock (0.5 mA, 2 second, scrambled current). During this 3-minute test, the mice received 1 shock during the last 2 seconds of a 30 second tone/light exposure. On Day 3, contextual conditioning (as determined by freezing behavior) was measured during a 5-minute test in the chamber where the mice were trained (context test). The next day, cued conditioning (CS+ test) was tested. The mice were placed in a novel context for 3 minutes, after which they were exposed to the conditioned stimuli (white light + tone) for 3 minutes. To create a novel context, the chamber was covered with new walls (white opaque plastic creating a circular compartment in contrast to a clear plastic square compartment) and a new floor (white opaque plastic in contrast to metal grid). Animals were returned to home cage after the testing period.

#### 2.3E Grip Strength

Mice were placed in the testing room 1 hour before starting and were subjected to standardized grip strength test. In brief, mice were placed on a TSE Systems grip strength meter that allowed for a 2-paw grip. Once mice gripped the meter, they were gently pulled by the base of the tail, parallel to the table surface, until they released their grip. The force exerted as the moment of release was recorded as the grip strength. Measurements were repeated three times with a one-minute rest interval.

### 2.4 Gene expression analysis

#### 2.4A Nanostring nCounter AD and neuroinflammation panels

Assays were performed as described previously [5]. In brief, 100 ng aliquots of hippocampal RNA were extracted using the PureLink RNA kit (Invitrogen, 12183018A) and run via the Nanostring nCounter analysis system (NanoString Technologies, Seattle, WA, USA) at the University of Alabama Nanostring Core. After codeset hybridization overnight, the samples were washed and immobilized to a cartridge using the NanoString nCounter Prep Station. Cartridges were scanned in the nCounter Digital Analyzer at 555 fields of view for the maximum level of sensitivity. Raw reads and the Nanostring Mm_AD30_v1.1 probe annotation codeset were imported into the Nanostring proprietary software, nSolver 4.0, and QC was performed following company protocol guidelines.

Counts for target genes were normalized to house-keeping genes included in the panel. Specifically, background correction was performed using the negative control at the cutoff of mean + 2 standard deviation. Housekeeping genes were used for normalization based on geometric mean. All p values were adjusted using a false discovery rate (FDR) correction of 1% for multiple comparisons. Differential expression and pathway scoring analysis, as well as graphical QC representation were performed in the nSolver Advanced Analysis Software 2.0 using a custom analysis to utilize generated normalized counts. Differential gene expression values were presented as 5xFAD vs. 5xFAD;cTFEB:HSACre hippocampal RNA expression counts. A p-value < 0.05 was considered significant. Volcano plots were generated with the ggplot2 package in RStudio [18] using significant genes and predetermined cutoff value. The pathway scoring module summarizes a single score from the data from a pathway’s genes.

### 2.5 Protein extraction

Protein lysates from skeletal muscle or cortex tissue from female control, 5xFAD or 5xFAD;cTFEB;HSACre mice were prepared as previously described [5, 7, 8] with a few modifications. Flash frozen perfused tissue (quadriceps and cortex) were added to a 2.0 mL plastic tube with silica beads (MP Biomedical, 116913100) and homogenized using a FastPrep-24 5G bead grinder and lysis system (MP Biomedical, 116005500) in radioimmunoprecipitation assay (RIPA) lysis and extraction buffer (Invitrogen, 89900), 1X Halt Protease and Phosphatase Inhibitor Cocktail (Invitrogen, 78442) and 1% (w/v) SDS. The supernatant was transferred to a new 1.5 mL plastic tube and centrifuged (Eppendorf, 5425R) at 1,500xg for 10 minutes at 4°C to remove debris. The resulting supernatant was transferred onto a new 1.5 mL plastic tube. Protein concentration was quantified using a Pierce^TM^ BCA Protein Assay (ThermoFisher Scientific, 237227) and frozen at -80°C until downstream analysis.

### 2.6 Synaptosome extraction

Total protein lysates and synaptosomes were extracted from 3-months-old C57BL/6 male mice following previously established protocols [19, 20] with a few modifications. In brief, flash frozen perfused cortex tissue was added to a 2.0 mL plastic tube with silica beads (MP Biomedical, 116913100) and homogenized using a FastPrep-24 5G bead grinder and lysis system (MP Biomedical, 116005500) in Buffer A, which consisted of 10 mM HEPES (Millipore Sigma, H3375) (pH 7.4), 2 mM EDTA (Millipore Sigma, E4884), 5 mM sodium orthovanadate (Millipore Sigma, S6508), 30 mM sodium fluoride (Millipore Sigma, 201154), 20 mM β-glycerol phosphate (Millipore Sigma, G5422), and protease inhibitor cocktail (Millipore Sigma, P2714), all diluted in 50 mL of water to a total volume of 50 mL. Tissue homogenates were then split onto two 1.5 mL plastic tubes.

To extract total protein lysates, 3% (v/v) Triton X-100 (Millipore Sigma, X100) was added to tissue homogenates and allowed to incubate in ice for 30 minutes with gentle shaking followed by centrifugation at 10,000xg for 15 minutes at 4°C. The resulting supernatant was transferred onto a new 1.5 mL Eppendorf tube; protein concentration was quantified using a Pierce^TM^ BCA Protein Assay (ThermoFisher Scientific, 237227) and frozen at -80°C until downstream analysis.

To extract synaptosomes, total homogenates were centrifuged for 500xg for 5 min at 4°C, and the resulting supernatant was separated from the nuclear pellet and transferred onto a new 1.5 mL tube followed by centrifugation at 10,000xg for 15 minutes at 4°C. The cytosolic supernatant was transferred onto a new 1.5 mL tube, while the pellet containing the crude synaptosome fraction was dissolved in 200 μL Buffer B, which consisted of 10 mM HEPES (pH 7.4), 2 mM EDTA, 2 mM EGTA (Millipore Sigma, E3889), 5 mM sodium orthovanadate, 30 mM sodium fluoride and 20 mM β-glycerol phosphate, 1% (v/v) Triton X-100 protease inhibitor cocktail, all diluted in 50 mL of water to a total volume of 50 mL. Protein concentration was quantified using a Pierce^TM^ BCA Protein Assay and frozen at -80°C until downstream analysis.

### 2.7 Immunoblot analysis

Western blot analyses were conducted as previously described [5]. In brief, 5-10 μg (synaptosome preps) or 20-40 μg (for total tissue) of homogenized protein lysates were mixed in appropriate volumes with NuPAGE LDS sample buffer (ThermoFisher Scientific, NP0007) and NuPAGE sample reducing reagent (ThermoFisher Scientific, NP0004) and were denatured at 70°C for 10 minutes. Samples were then loaded per lane onto Any kDa, 4-15% or 10% Mini-PROTEAN TGX Stain-Free Precast Gels (BioRad, 4568123, 4568083 or 4568033). Gels were activated by exposure to UV light for 45 seconds using a Chemidoc (BioRad, 12003153), and protein lanes were transferred onto a 0.45 μm PVDF membrane (BioRad, 1704275) using the BioRad Turbo Transfer system (BioRad, 1704150). Post-transfer membranes were blocked in 5% (w/v) BSA in TBS + 0.1% (v/v) Tween-20 (TBST) or 5% (w/v) skimmed milk in TBST at RT for 1 hour. Membranes blocked with milk were washed three times with TBST to remove excess milk prior to incubation with primary antibodies. Skeletal muscle membranes were incubated with anti-FLAG or PSAP antibodies. Cortex membranes were incubated with anti-brain derived neurotrophic factor (BDNF), neurotrophin 4 (NTF4), synaptosomal-associated protein 25 (SNAP25), synaptophysin 1 (SYP1), synaptotagmin 1 (SYT1), synapsin 1 (SYN1), post-synaptic density protein (PSD95), synaptosome associated protein 97 (SAP97), SAP102, glutamate ionotropic receptor AMPA type subunit 1 (GluA), actin β, PSAP/saposin-C or β tubulin 3. See **Table S1** for full details. Finally, all membranes were subjected to gentle shaking at 4°C overnight, washed three times with 1X TBST and incubated with appropriate secondary antibodies for 1-2 hours with gentle shaking at RT. Membranes were washed twice with TBST and once with TBS, and imaged using the Chemidoc (BioRad, 12003153). All data were analyzed using densitometry analysis normalized to an appropriate loading control (Total StainFree, actin β or tubulin depending on lysate origin) in ImageLab.

### 2.8 Prosaposin ELISA

Blood was collected from mice via cardiac puncture and plasma separated using standard coagulation/centrifugation protocols as described above. Collected plasma was diluted 1:2222 and processed through a mouse PSAP ELISA kit (Abcam, ab277399) following the manufacturer’s protocol. In brief, 100 μL of standard calibrator or sample was added to a plate and allowed to incubate for 2.5 hours with gentle shaking at RT. Wells were washed 4X with wash buffer and 100 μL of biotinylated anti-PSAP antibody was added to each well and allowed to incubate for 1 hour with gentle shaking at RT. Wells were then washed four times with wash buffer and a 100 μL of HRP-streptavidin concentrate was added to each well and allowed to incubate for 45 minutes with gentle shaking at RT. After four washes, 100 μL of TMB substrate reagent was added to each well and allowed to incubate for 30 minutes at RT. Finally, 50 μL of stop solution was added to each well and the plate was read immediately at 450 nm (Tecan, 30190085). The concentration of PSAP was determined using a standard serially diluted curve of known concentration.

### 2.9 PS18 Administration

An 18-mer derived peptide (LSELIINNATEELLIKGL; PS18) from the amino-terminal hydrophilic sequence of the rat saposin-C domain was obtained from Phoenix Pharmaceuticals (324-431), dissolved in dimethyl sulfoxide (DMSO), which was diluted in sterile normal saline (0.9%) into a solution with 1% DMSO (v/v) the day before injections. Each mouse was subcutaneously injected with a final concentration of 2.0 mg/kg PS18 or 1% DMSO alone once a day for seven days. Mice were sacrificed on the 8th day and tissue was collected and processed as described above.

## 3. RESULTS

### 3.1 Conditional skeletal muscle TFEB overexpression reduces Aβ accumulation and rescues neurocognitive behavioral deficits 5xFAD female mice

Previous work from our lab has shown that skeletal muscle-TFEB overexpression is sufficient to improve learning and memory and reduce neuroinflammation and CNS protein aggregation during healthy aging and in the MAPT P301S mouse model of tauopathy [5]. To investigate whether muscle-TFEB overexpression is also neuroprotective in the context of another primary neuropathological hallmark of AD (i.e., Aβ), we generated 5xFAD;cTFEB;HSACre triple transgenic mice to achieve muscle-TFEB overexpression in the well-characterized plaque-toxicity 5xFAD transgenic background [21] (**Figure 1A**). For this work, we focused our intervention on 5xFAD transgenic female mice, as they exhibit an earlier and more severe phenotypic presentation of AD-associated pathology than their male counterparts [22], including more severe behavioral deficits, a stronger neuroinflammatory response associated with Aβ plaque deposition and other metabolic deficits. These observations also align with the known increased prevalence and risk for AD in human females compared to males [23].

We first confirmed robust skeletal muscle 3x-FLAG-TFEB overexpression via immunoblotting in skeletal muscle lysates of 5xFAD;TFEB;HSACre, but not 5xFAD;TFEB or 5xFAD;HSACre littermates (hereafter referred to as 5xFAD, **Figure 1B**). There were no significant changes in the levels of TFEB protein expression between 4 months (early symptomatic) and 8 months of age (late symptomatic) in 5xFAD;cTFEB;HSACre skeletal muscle (**Figure S1A**). Importantly, we also saw no detectable exogenous expression of TFEB in the CNS of 5xFAD;TFEB;HSACre mice via immunoblotting of cortical lysates from the same cohorts (**Figure 1B**). This confirms inducible TFEB expression at stable levels in skeletal muscle across two stages of disease progression in this model (**Figure 1B**), similar to what we observed in our MAPT P301S;cTFEB;HSACre model [5].

Given the profound decreases in the accumulation of neurotoxic hyperphosphorylated tau in our MAPT P301S work [5], we first examined whether overexpression of TFEB in skeletal muscle modifies the accumulation of Aβ in the CNS of 5xFAD mice. Consistent with previous work deep-phenotyping the progressive age-associated increase in plaque burden in this model [24], the 6E10+ fraction of the 5xFAD cortex and hippocampus increased from around 1% at 4 months of age up to ∼8-10% at 8 months of age. Interestingly, we detected significant reductions in the 6E10-positive fraction area in the 5xFAD;cTFEB;HSACre cortex (**Figure 1C-D, Figure S1B**) and hippocampus (**Figure 1E-F, Figure S1C**) at both ages compared to their 5xFAD littermates. This suggests that muscle-TFEB overexpression reduces Aβ-peptide accumulation and/or aggregation in disease-relevant CNS regions of 5xFAD;cTFEB;HSACre transgenic mice.

We have previously reported that muscle-TFEB overexpression improves cognitive performance in aged mice compared to age-matched controls [5]. Given the significant reductions in plaque burden observed in the CNS of 5xFAD;cTFEB;HSACre mice at 8 months of age (**Figure 1C-F**), we next interrogated behavioral performance in our cohorts at this symptomatic stage. As previously reported [14, 32], 5xFAD transgenic mice weighed less than their control littermates at 8 months of age, a phenotype believed to reflect on-going metabolic imbalance (**Figure 1G**). Interestingly, the body weight of age-matched 5xFAD;cTFEB;HSACre mice was not significantly different than their control littermates (**Figure 1G**), consistent with our previous findings that muscle-TFEB overexpression improves overall skeletal muscle and metabolic health [5, 25]. We found no differences among any of groups on grip strength tests (**Figure 1H**), consistent with mild to no muscle weakness in this model at this age [24].

We next evaluated neurocognitive performance in our cohorts using a battery of tests with well-characterized deficits for the 5xFAD model, beginning with nest building to assess general behavior, well-being, and sensorimotor gating [26]. While control animals scored high on a 24- hour nest building test (average score of 2.1, maximum score of 3), 5xFAD transgenic mice scored quite poorly (average score of 0.4), rarely exhibiting interest in the provided nestlet (**Figure 1I**). Conversely, 5xFAD;cTFEB;HSACre mice had significantly higher nesting scores (average score of 1.4), displaying increased interactions with the nestlets and building more complex nests (**Figure 1I**). This score was not significantly different than the one observed in control mice.

Female 5xFAD transgenic mice have well-characterized deficits in spatial learning and memory in the Barnes maze [24, 27, 28], spending less time in the target quadrant (**Figure 1J**) and a longer latency to the target hole (**Figure 1K**) than their control littermates. Interestingly, 5xFAD;cTFEB;HSACre mice performed at the same level as control animals in both metrics (**Figure 1J and K**), spending more time in the target zone and escaping the maze faster than their 5xFAD counterparts, without any significant differences in motor activity among any our cohorts (**Figure S2A-C)**. Finally, 5xFAD;cTFEB;HSACre mice performed equally as well as their control littermates and significantly better than their 5xFAD littermates when we assessed associative learning memory via fear conditioning testing (**Figure 1L**). Collectively, this battery of neurocognitive behavioral tests suggests that muscle-TFEB overexpression rescues neurocognitive deficits in decision-making, spatial memory and associative learning in 8- months-old 5xFAD mice.

### 3.2 Skeletal muscle TFEB overexpression does not alter neuroinflammation markers in the CNS of 5xFAD female mice

Neuroinflammation, the chronic activation of glial cell toward pro-inflammatory phenotypes, is a well-characterized pathological hallmark of AD [29] that is recapitulated in the brains of 5xFAD mice [30]. Given the robust reductions in morphological and transcriptional markers of neuroinflammation we observed in response to skeletal muscle TFEB overexpression in the context of tau toxicity [5], we wondered whether neuroinflammation was once again a target of skeletal muscle-TFEB-dependent neurotrophic signaling in 5xFAD;cTFEB;HSACre mice. To assess glial activation in situ, we first examined astroglial morphology in the dentate gyrus of the hippocampus (**Figure 2A and S3A**). As expected, and reflecting the high degree of neuroinflammation of this model at symptomatic stages of disease, we detected a large age-dependent increase in the GFAP-positive (astrocytes) and IBA1-positive (microglial) fractions of the hippocampus in 5xFAD transgenic mice compared to their control littermates (**Figure 2B and S3B**). We also confirmed significant increases in morphological markers of pro-inflammatory states, including the volume of hippocampal IBA1+ microglia (**Figure 2C**) and the number of GFAP+ astrocytes (**Figure 2D**) in the 5xFAD hippocampus relative to their control littermates. However, and in opposition to what we observed in our MAPT P301S;cTFEB;HSACre model [5], the fraction area coverage, microglial volume or astrocyte number in the hippocampus of 5xFAD;cTFEB;HSACre mice was comparable to that observed in 5xFAD mice at either age examined (**Figure 2A-D and S3A-D)**.

**FIGURE 2.**
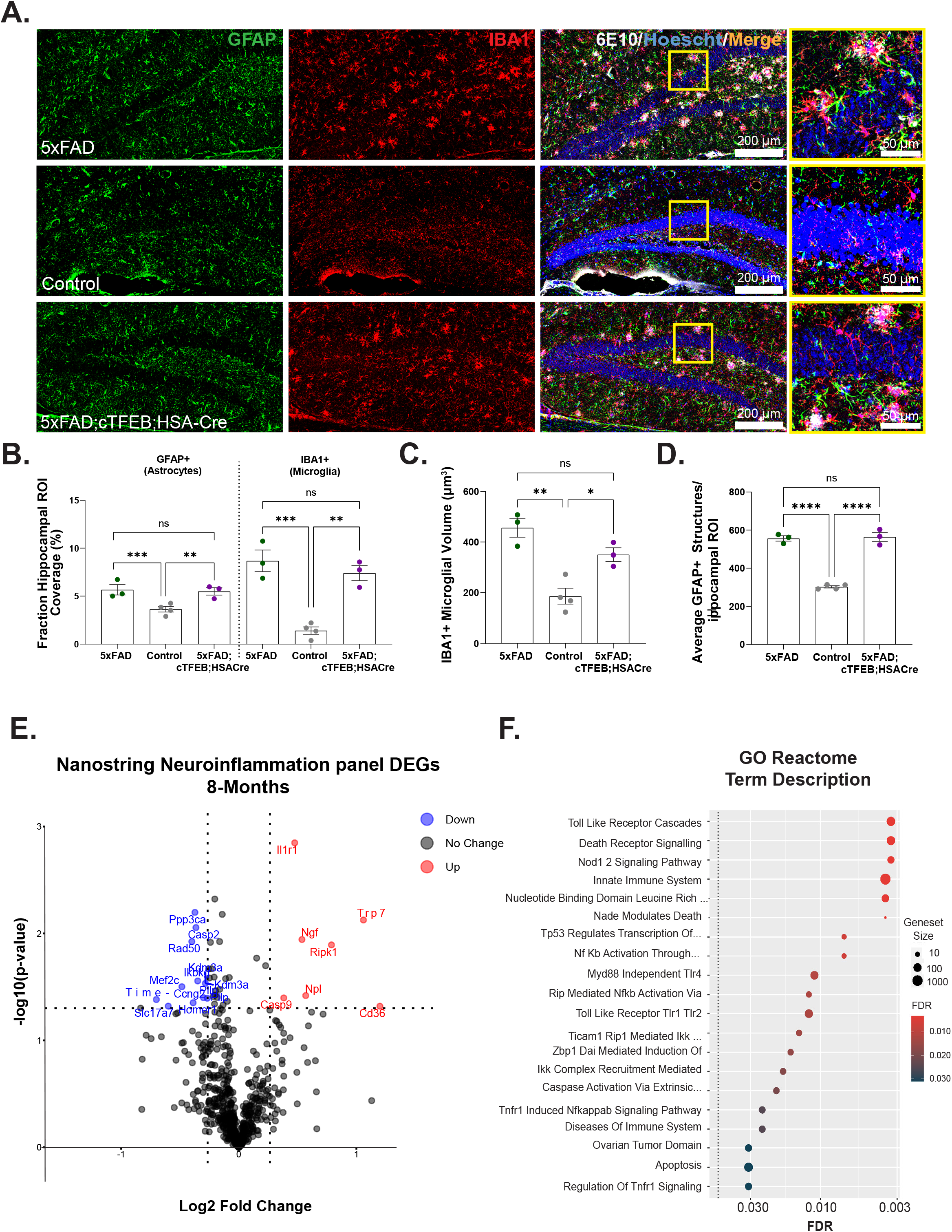
Skeletal muscle-TFEB overexpression did not alter neuroinflammation in 8- months-old 5xFAD transgenic female mice. **(A)** Representative images of the dentate gyrus of the hippocampus stained for astrocytes (GFAP, green) microglia (IBA1, red), 6E10 (white) and Hoechst (blue). Scale bars as shown. **(B)** Quantification of astrocyte and microglia load. **(C)** Volume of microglia. **(D)** Number of GFAP+ astrocytes. **(E)** Volcano plot of differentially expressed hippocampal transcripts quantified using a Nanostring Neuroinflammation panel comparing 5xFAD;cTFEB;HSACre vs. 5xFAD 8-months-old female transgenic mice. Up-regulated transcripts in red, down-regulated transcripts in blue. **(F)** Dot plots of the gene ontology (GO) enrichment analysis according to a reactome panel. Statistical comparison was performed using one-way ANOVA and post hoc multiple comparisons, ^∗^p < 0.05, ^∗∗^p < 0.01, ^∗∗∗^p < 0.001 , n.s. non-significant. Data is represented as mean ± SEM.

Targeted transcriptional analysis in hippocampal RNA lysates using the Nanostring Neuroinflammation panel revealed 11 down-regulated and 7 up-regulated genes in the 5xFAD;cTFEB;HSACre hippocampus relative to their 5xFAD counterparts (**Figure 2E**). None of the 24 Nanostring functional pathway scores in this panel were significantly different among groups (not shown). GO biological process analysis of the differentially expressed genes identified mostly unrelated signaling pathways, including response to inorganic substance, positive regulation of molecular function and cellular response to stress (**Figure S3E**). Top Reactome terms included some immune-associated pathways like toll-like receptor cascades, death receptor signaling and innate immune system (**Figure 2F**). We also confirmed no differences in expression levels of classical neuroinflammatory markers *Ccl2, Il6* or *Nfkβ* in hippocampal RNA lysates (not shown).

Altogether, our morphological and transcriptional analyses indicate minimal changes in neuroinflammatory states in the hippocampus of 5xFAD;cTFEB;HSACre mice relative to their 5xFAD littermates, despite robust reductions in Aβ accumulation (**Figure 1C-F**). Furthermore, it indicates that the observed behavioral neurocognitive improvements in 5xFAD;cTFEB;HSACre mice (**Figure 1I-L**) likely occurs through a mechanism independent of neuroinflammatory associated pathways.

### 3.3 Skeletal muscle TFEB overexpression promotes transcriptional remodeling associated with synaptic function in the CNS of 5xFAD female mice

Given the mild transcriptional effects observed with the Nanostring Neuroinflammation panel in 5xFAD;cTFEB;HSACre hippocampi (**Figure 2E**), we next proceeded to directly interrogate AD-associated signaling using the Nanostring Alzheimer’s Disease panel. Importantly, this panel was built utilizing the 5xFAD model in addition to human samples, creating a much more targeted disease profile [31–33]. We identified 15 differentially expressed genes in 5xFAD;cTFEB;HSACre hippocampal RNA lysates when compared to their 5xFAD littermate controls (**Figure 3A**). Of these, 12 were down-regulated and only 3 were up-regulated, reflecting the high degree of disease-association built into the panel footprint [31–33]. Two of these genes were associated with protein aggregation (*Kctd1*, potassium channel tetramerization domain containing 1) [34] and Aβ amyloidosis formation (*Phyhip*, phytanoyl-CoA 2-hydroxylase interacting protein) [35], consistent with our findings of reduced plaque abundance in the CNS of 5xFAD;HSACre mice (**Figure 1C-F**). Strikingly, many of the other down-regulated genes coded for multiple hits on the synaptic regulatory network, including, *Stxbp5l* (a syntaxin-binding protein), protein kinase C related genes *Prkca* and *Prkcb*, *Arhgef9* and *Mef2c.* Additional significantly down-regulated genes that were associated with different aspects of synaptic signaling were *Kcng5* (potassium voltage-gated channel subfamily J member 5), *Chrm1 (*acetylcholine receptor), *Pde1a* (phosphodiesterase-1) and *Cnler2/ Cnksr2* (connector enhancer of kinase suppressor of Ras 2). Consistent with transcriptional targeting of the synaptic signaling network, we also detected significant up-regulation of neurotrophic-associated *Gss* (glutathione synthetase), as well as *Cbln1* (precerebellin), another promoter of synaptic plasticity [36] (**Figure 3A**). Analysis of Nanostring functional pathway scores also confirmed profound rescue of gene signature scores for vesicle trafficking (**Figure 3B**), neuronal connectivity (**Figure 3C**) and chromatin modification (**Figure 3D**), all of which are pathways known to be dysregulated in symptomatic 5xFAD female mice [22, 37]. Interestingly, we also observed non-significant trends for rescue of the lipid metabolism signature gene cluster (**Figure 3E**), another classical pathological hallmark of AD [38].

**FIGURE 3.**
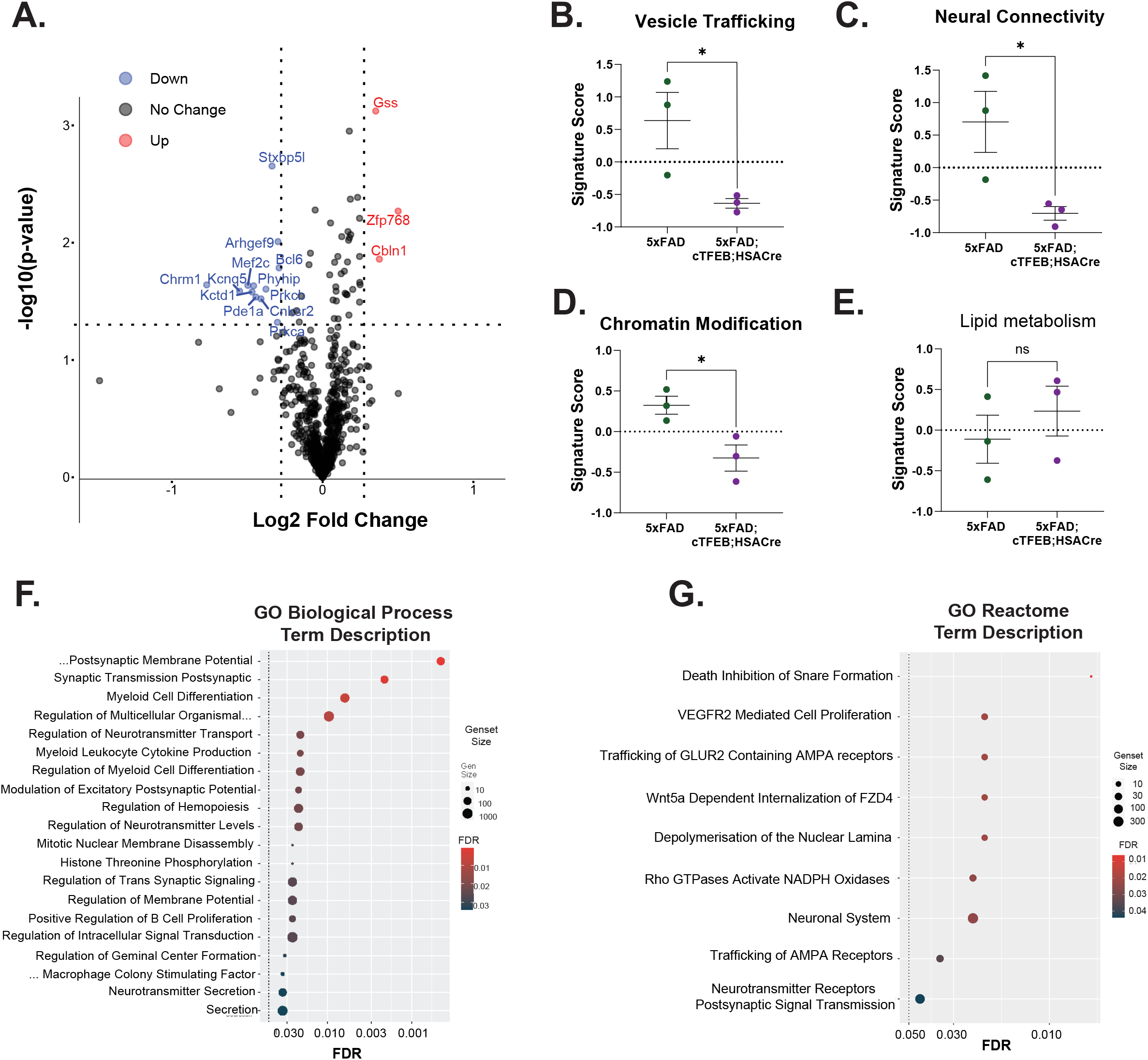
Skeletal muscle-TFEB overexpression alters synaptic-associated gene transcriptional expression in 5xFAD 8-month-old transgenic female mice. **(A)** Volcano plot of differentially expressed hippocampal transcripts quantified using a Nanostring AD panel for comparing 5xFAD;cTFEB;HSACre vs. 5xFAD 8-months-old transgenic female mice. Up-regulated transcripts in red, down-regulated transcripts in blue. (**B-E)** Associated signature score for pathways involving vesicle trafficking **(B**), neural connectivity **(C)**, chromatin modification **(D)** and lipid metabolism **(E).** Gene scores are shown relative to control (5xFAD-) group (dotted line). **(F-G)** Dot plots of the gene ontology (GO) enrichment analysis for transcripts involved in biological processes **(F)** or according to a reactome panel **(G)**. Statistical comparison was performed using an independent t-test, ^∗^p < 0.05, ^∗∗^p < 0.01, ^∗∗∗^p < 0.001, n.s. non-significant. Data is represented as mean ± SEM.

We next performed gene set enrichment analysis on the DEGs in the 5xFAD;cTFEB;HSACre hippocampi which confirmed significant changes on gene pathways associated with pre- and post-synaptic membrane potential and transmission pathways (GO Biological Processes, **Figure 3F**), and trafficking of GLUR2 containing AMPA receptors and VEGFR2 mediated signaling (GO Reactome, **Figure 3G**). Altogether, this transcriptional analysis strongly suggests that muscle-TFEB overexpression transcriptionally promotes remodeling of the synaptic networks in 5xFAD female transgenic mice.

### 3.4 Skeletal muscle TFEB overexpression increases levels of synaptic proteins in the CNS of 5xFAD female mice

To validate whether these transcriptional changes associated with synaptic function (**Figure 3**) could underlie the behavioral improvements we observed in 5xFAD;cTFEB;HSACre transgenic mice (**Figure 1I-L**), we assayed levels of synaptic proteins in cortical lysates of all our groups (**Figure 4**). We first confirmed previously reported deficits in levels of neurotrophic factors such as brain derived neurotrophic factor (BDNF, particularly of the cleaved, secreted form), and neurotrophin 4 (NTF4) (**Figure 4A**) in the cortex 8-months-old 5xFAD transgenic mice [39–42]. Interestingly, while the pre-synaptic protein markers SNAP25, SYP1 and SYT1 were unchanged, post-synaptic markers PSD95, SAP102 and SAP97 showed significant reductions in 5xFAD cortical lysates at this age (**Figure 4B**). These markers were not altered in 5xFAD 4-months-old mice (**Figure S4A-C**), confirming their reflection of neuropathological processes occurring in post-symptomatic stages in this model. Strikingly, protein levels of most of these pre- and post-synaptic markers were significantly increased in the 5xFAD;cTFEB;HSACre cortex at 8 months of age, even to levels above their littermate controls (**Figure 4A-B**). Overall, this data strongly implicates improvements in synaptic function, likely driven by increases in neurotrophic signaling, as a potential mechanism underlying the neuroprotective benefits observed in 5xFAD;TFEB;HSACre transgenic mice.

**FIGURE 4.**
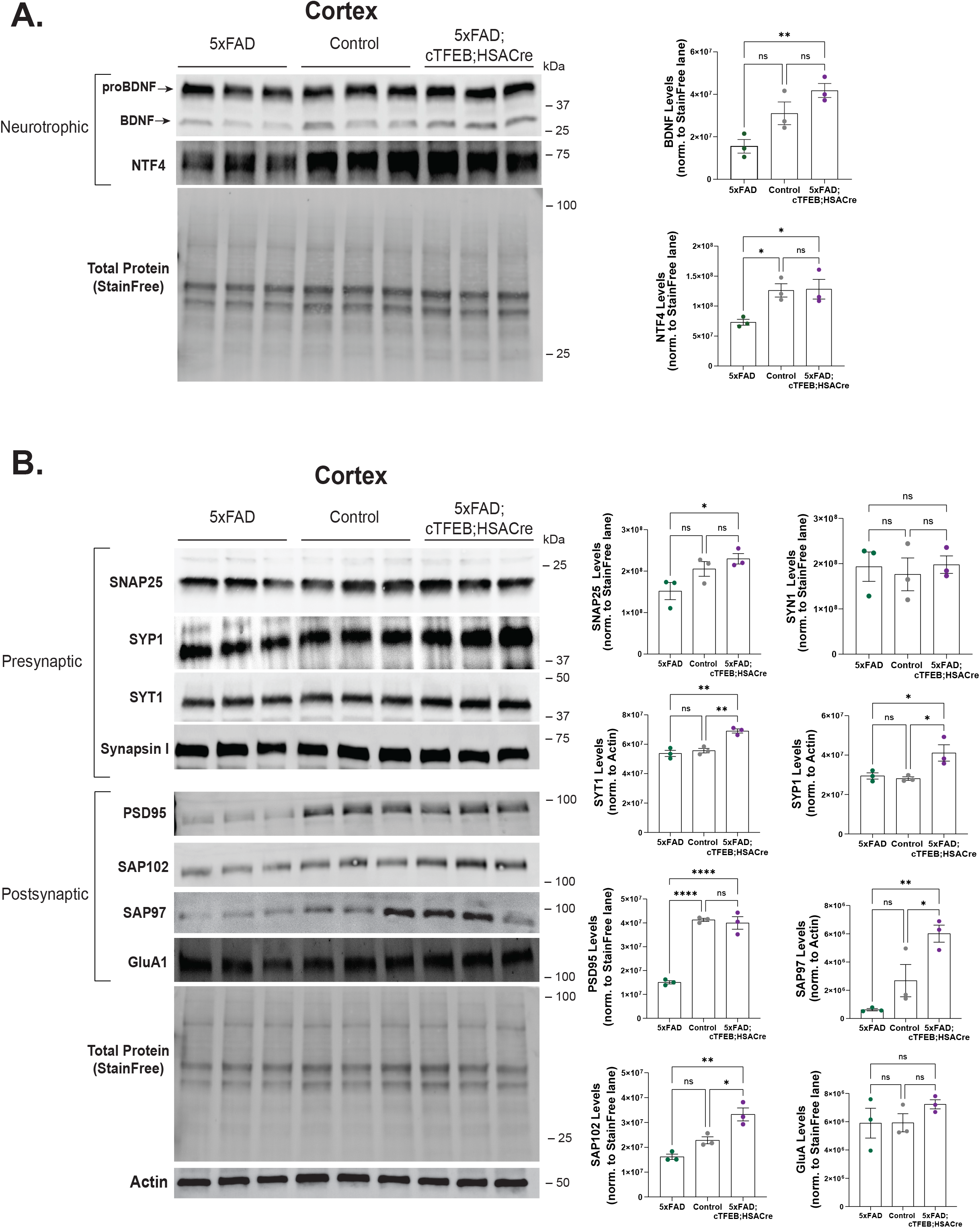
Skeletal muscle-TFEB overexpression enhances neurotrophic signaling and synaptic integrity in brains of 5xFAD 8-months-old transgenic female mice. **(A-B)** Immunoblot for several neurotrophic **(A)**, pre- and post-synaptic **(B)** markers using 8-months-old mice cortical protein lysates. Total StainFree protein or actin β was used a loading control. Densitometry quantification for brain-derived neurotrophic factor (BDNF), neurotrophin 4 (NTF4), synaptosome-associated protein 25 (SNAP25), synaptophysin I (SYP1), synaptotagmin I (SYT1) synapsin I (SYN1), postsynaptic density protein 95 (PSD95), synapse-associated proteins 97 (SAP97) and 102 (SAP102) and glutamate ionotropic AMPA receptor subunit 1 (GluA1) normalized to either Total StainFree for each protein lane (for loadings of 40-50 ug of total protein) or actin β band (for loadings of 20-25 ug of total protein) densitometry, as shown on graphs on the right. Statistical comparison was performed using one-way ANOVA and post hoc multiple comparisons, ^∗^p < 0.05, ^∗∗^p < 0.01, ^∗∗∗^p < 0.001, n.s. non-significant. Data is represented as mean ± SEM.

### 3.5 Skeletal muscle-targeted TFEB overexpression increases release of CNS-targeting myokine Prosaposin in 5xFAD;cTFEB;HSACre transgenic mice

We have previously demonstrated that muscle-TFEB overexpression is a powerful activator of the muscle-to-brain axis [5]. We further identified several potentially secreted proteins in TFEB-overexpressing skeletal muscle as potential new myokines with CNS targeting effects, including the precursor to lysosomal sphingolipid hydrolase activator proteins known as prosaposin (PSAP) [5]. Various studies have recently highlighted a potential neuroprotective effect for PSAP, whether as the full-length protein, saposin C (one of its biologically active cleavage products) or a saposin C-derived 18-mer derived peptide (PS18). Indeed, PSAP and its derivatives prevent neuronal degeneration and axonal loss [43, 44], stimulate nerve regeneration [45], neuronal growth [46], and neuroprotection in neurodegenerative model systems [11, 47–49]. Building on this body of evidence, we posited that PSAP may act as pivotal myokine facilitating muscle-to-brain crosstalk in our 5xFAD;cTFEB;HSACre mice.

In agreement with this, we confirmed that as predicted by our previous proteomics studies in young and aged cTFEB;HSACre transgenic mice [5], total PSAP protein levels were twice as high in 8-months-old 5xFAD;cTFEB;HSACre skeletal muscle relative to 5xFAD transgenic muscle (**Figure 5A**). Interestingly, we also detected a milder but still significant increase in PSAP and cleaved isoform saposin C in the cortex of the same mice (**Figure 5B**), suggesting systemic elevations of PSAP levels in 5xFAD;cTFEB;HSACre transgenic mice. Consistent with this hypothesis, ELISA testing confirmed large increases in circulating PSAP protein in 5xFAD;cTFEB;HSACre plasma (∼200,000 pg/ml) compared to either control or 5xFAD plasma (∼100,000 pg/ml) (**Figure 5C**), strongly suggesting that skeletal muscle PSAP overproduction via TFEB overexpression results in increased PSAP secretion into circulation.

**FIGURE 5.**
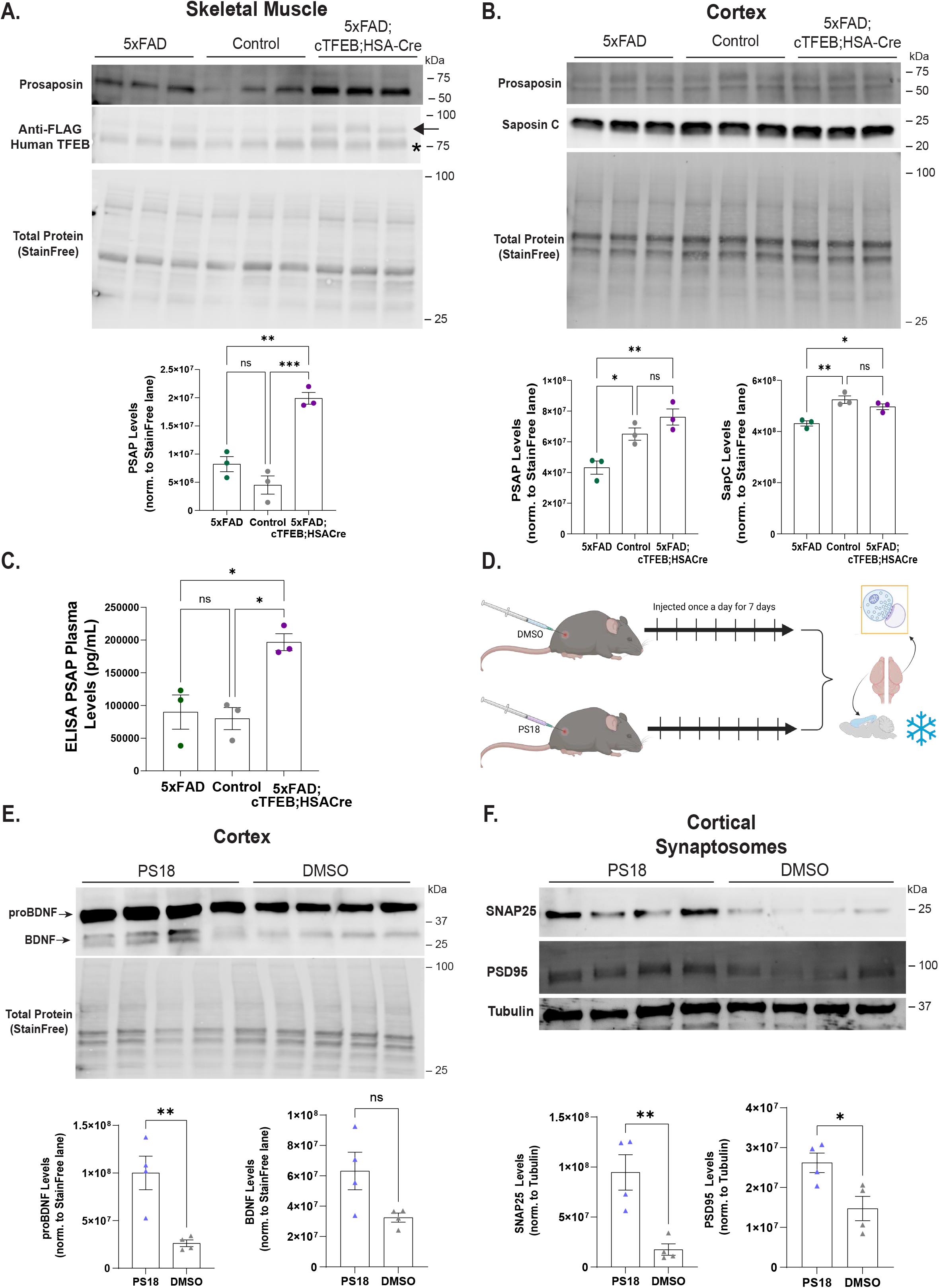
Prosaposin (PSAP) and PS18 as a novel myokine enriched in 5xFAD;cTFEB;HSACre mice that recapitulates enhancement of neurotrophic signaling and synaptic integrity. **(A-B)** Immunoblots for PSAP and 3x-FLAG in skeletal muscle **(A)** and for PSAP and saposin-C in cortex (**B**). Transgenic 3x-FLAG-TFEB protein weighs ∼75 kDa. Arrow highlights TFEB protein, asterisk (*) highlights a non-specific band. Densitometry quantification normalized by Total StainFree protein lane densitometry is shown below. **(C)** Enzyme-linked immunosorbent assay (ELISA) for PSAP in peripheral plasma of 8-months-old female mice. **(D)** Schematic illustration of PS18 or dimethyl sulfoxide (DMSO) subcutaneous injection in healthy 3-months-old C57/B6L male mice and subsequent isolation of cortical protein lysates and synaptosome-enriched fractions. **(E)** Immunoblot for BDNF in cortical protein lysates from PS18- or DMSO-injected C57/B6L male mice. Densitometry quantification for proBDNF and BDNF normalized by Total StainFree protein lane densitometry is shown below. **(F)** Immunoblots for SNAP25 and PSD95 in cortical synaptosome-enriched fractions from PS18- or DMSO-injected C57/B6L male mice. Densitometry quantification normalized to β tubulin is shown below. Statistical comparison was performed using one-way ANOVA and post hoc multiple comparisons or independent t-tests, ∗p < 0.05, ∗∗p < 0.01, ∗∗∗p < 0.001, n.s. non-significant. Data is represented as mean ± SEM.

Multiple studies have demonstrated that supplementation of individual myokines in the periphery results in potent neurotrophic effects in the CNS [50–53]. To test the potential role of PSAP as a novel myokine mediating the synaptic remodeling observed in the CNS of 5xFAD;cTFEB;HSACre mice, we subcutaneously injected 3-months-old C57B6/L male mice once a day for 7 days with PS18 (**Figure 5D**), the 18-mer derived peptide from the amino-terminal hydrophilic sequence of the rat saposin C-domain. This PS18 injection paradigm has previously been shown to effectively target the CNS, reducing Aβ_1–42_ neurotoxicity and memory deficits [54]. Consistent with our hypothesis that peripheral PSAP-signaling mediates the synaptic remodeling observed in the 5xFAD;cTFEB;HSACre CNS, immunoblot analysis of cortical lysates revealed a robust increase in both pro- and active BDNF isoforms in PS18-injected animals relative to their DMSO injected controls (**Figure 5E**). To obtain direct insights into changes localized to the synaptic compartment, we next isolated cortical synaptosome fractions enriched in pre- and post-synaptic machinery from the cortex of PS18 and DMSO-injected mice. Immunoblot analyses of cortical synaptosomes showed that peripheral PS18-injection increased the abundance of pre-synaptic protein SNAP25 and post-synaptic protein PSD95 (**Figure 5F**) as well as trends towards increases for SYP1, SYT1, and SAP97 in cortical synaptosome-enriched fractions from PS18-injected animals (**Figure S5B**), similar to what we observed in 5xFAD;cTFEB;HSACre cortices (**Figure 4**). Altogether, these provocative data suggest that peripheral supplementation of PS18 is sufficient to activate neurotrophic signaling and promote synapse remodeling in the CNS. Furthermore, it implies that the neurocognitive benefits seen in 5xFAD;cTFEB;HSACre mice can be attributed, at least in part, to increased secretion of muscle-originating PSAP into circulation as a novel CNS-targeting myokine.

## 4. DISCUSSION

Over the last decade, growing evidence has suggested that the periphery contributes to the etiology of neurodegenerative diseases, including AD [5, 12, 13, 53, 55–57]. In particular, activation of the muscle-to-brain axis and its associated myokines collectively target central markers of CNS dysfunction, decreasing neuroinflammation and reducing proteotoxicity in the CNS, all key factors in AD pathogenesis [23]. Indeed, peripheral delivery of CNS-targeting circulating factors such as Fncd5/irisin [53], platelet factor-4 (Pf4) [58] and glycosylphosphatidylinositol-specific phospholipase D1 (Gpld1) [59] all individually promote significant remodeling of the mouse brain, although the precise mechanisms driving these benefits remain unknown. Here we provide direct evidence demonstrating benefits in the CNS after enhancing skeletal muscle TFEB signaling in the context of Aβ-induced toxicity, and validate PSAP as a new myokine with CNS-targeting effects.

Similar to what we previously reported in a model of tau-associated toxicity [5], we discovered that overexpression of TFEB in skeletal muscle (**Figure 1B**) significantly improves multiple disease-associated hallmarks in 5xFAD female transgenic mice. Indeed, we confirmed robust reductions in plaque accumulation (**Figure 1C-F**) and profound synaptic remodeling (**Figure 3 and 4**) in the hippocampus and cortex of 5xFAD;cTFEB;HSACre transgenic mice. This is consistent with our previous work showing that expression of many genes related to synaptic transmission and plasticity are transcriptionally remodeled in the hippocampus of young and aged female cTFEB;HSACre transgenic mice, even in the absence of AD-associated pathologies [5]. Importantly, we confirm that these biochemical findings translate into a significant rescue on behavioral performance across a battery of neurocognitive tests (**Figure 1G-L**) in symptomatic 5xFAD mice with skeletal muscle-TFEB overexpression. These results thus imply, for the first time, that activation of the muscle-to-brain axis via skeletal muscle-TFEB overexpression is a powerful augmenter of neurotrophic signaling and synaptic network remodeler in the CNS.

One of most interesting findings from this study was the mild effect of muscle-TFEB overexpression on markers of neuroinflammation (**Figure 2 and S3**). Whereas we observed major reductions in both morphometric and transcriptional markers of neuroinflammation in the dentate gyrus of the hippocampus of our MAPT P301S;cTFEB;HSACre mice [5], we failed to detect similar changes in the same brain region of 5xFAD;cTFEB;HSACre mice. We did detect some modest transcriptional expression differences in the 5xFAD;cTFEB;HSACre hippocampus using the targeted Nanostring neuroinflammation panel (**Figure 2E-F**), raising the possibility that these non-canonical inflammatory signaling pathways may not translate into detectable morphological changes in astrocytes and microglia at the ages examined. Indeed, our previous results in MAPT P301S mice displayed a sex-dimorphic neuroinflammatory rescue associated with muscle-TFEB overexpression. This suggests that in our current study, the protective effects on neuroinflammation in 5xFAD;cTFEB;HSACre female mice may be concealed due to a differential inflammatory baseline derived from sex. Interestingly, we have previously reported a similar uncoupling between mild modifications of neuroinflammation markers and robust neurocognitive improvements in our healthy aged cTFEB;HSACre without any AD-associated transgenes [5]. This suggests that skeletal muscle TFEB overexpression may promote differential muscle-to-brain axis signaling in a CNS-pathology and/or sex-specific manner, although this hypothesis remains untested.

PSAP (also known as sulfated glycoprotein 1, SGP-1) is a complex multifunctional protein, playing roles both intracellularly (as a regulator of lysosomal function) and extracellularly as a secreted factor [44]. The extracellular presence of full-length prosaposin suggests a discrete function for full-length prosaposin beyond its role as the saposin precursor protein. Indeed, full-length PSAP was recently found to be enriched in skeletal muscle exudates [60]. Furthermore, expression levels of PSAP are significantly increased in both skeletal muscle and plasma of exercised (running) rats [61], suggesting PSAP secretion is responsive to skeletal muscle-targeting interventions. In agreement with this, and using unbiased proteomics analysis, we had previously also predicted PSAP as one of the most highly upregulated secreted proteins in cTFEB;HSACre skeletal muscle [5]. Here, we validate our previous findings, and extend them to our new disease model, showing significant increases in the PSAP protein locally in skeletal muscle (**Figure 5A**) and systemically in circulation (plasma; **Figure 5C**) of 5xFAD;cTFEB;HSACre transgenic mice. Furthermore, we also demonstrate significant enhancement of neurotrophic and pre- and post-synaptic markers in the CNS after peripheral injections of a prosaposin-derived 18-mer peptide (PS18). Of note, whereas we report increases in BDNF protein levels in 5xFAD;cTFEB;HSACre (Figure 4A) and PS18-injected whole cortical lysates (**Figure 5E**), we only detected changes in synaptic protein markers in synaptosome-enriched fractions from PS18-injected mouse cortices (**Figure 5F** and **S5B**). We believe this reflects the effects of chronic increased circulating PSAP on the CNS in our 5xFAD;cTFEB;HSACre mice vs. the acute activation of PSAP-signaling associated with peripheral PS18 delivery. Indeed, although endogenous PSAP is enriched in the brain and has a well-known history of exhibiting neuroprotective actions [44], our work shows for the first time a role for muscle-originating peripheral PSAP in mediating CNS-targeting neuroprotective effects.

Two central gaps in knowledge in our current understanding of CNS-targeting myokines currently remain. First, the mechanism of transport of these myokines through circulation is a highly debated topic in the field. Many studies have identified myokines as part of the cargo contained within extracellular vesicles (EVs), including those originating from skeletal muscle [62]. However, delivery of recombinant proteins and/or AAV-packaged versions of myokines into peripheral circulation has also successfully targeted the CNS, suggesting that vesicle-bound entities may not be necessary for activation of the muscle-to-brain axis. Importantly, the content and abundance of myokines in circulation is highly responsive to exercise [53, 62–64], suggesting they may act as direct modulators of muscle-targeting interventions with known neurotrophic effects [5, 65]. Second, is the ability (and necessity) of myokines to cross the blood-brain barrier (BBB). While some work supports the potential transmembrane transport, diffusion or signal-transduction of these myokines across the blood-brain-barrier [53, 64], to date, the mechanisms that transmit the peripheral signal into the CNS remain unknown. Full length PSAP is unlikely to cross the BBB, so it may be instead carried within EVs as they are known to traverse across BBB [66–68] or it could be enzymatically digested to more soluble forms such as saposin C or PS18. Either way, these cleavage products of PSAP have come to be collectively known as ‘prosapeptides’ due to their reported neurotrophic effects. Additional research is needed to clarify these on-going questions.

Overall, this work connects, for the first-time, skeletal muscle TFEB overexpression to increased secretion of PSAP into circulation, and validate PSAP as a novel myokine with CNS-targeting effects in an Aβ-induced toxicity AD model (i.e., 5xFAD). Ultimately, this work reveals a potential new mechanism of muscle-to-brain communication during neurodegenerative disease through neuroprotective muscle-originating secreted protein signatures, ‘reuniting the body from the neck down’ [69].

## Acknowledgement

We thank all members of the Cortes lab past and present that contributed their support to this work. We also thank C. Finch for providing us with necessary reagents and intellectual discussion throughout. Authors have nothing to disclose.

