## Supplementary material for "Activation of the muscle-to-brain axis ameliorates neurocognitive deficits in an Alzheimer’s disease mouse model via enhancing neurotrophic and synaptic signaling": Table S1

**Table S1.** Detailed information for antibodies used in immunofluorescence and immunoblotting experiments.

| **Antibody** | **Company and Catalog #** | **Dilution** |
|---|---|---|
| **Immunohistochemistry** | | |
| Chicken anti-GFAP | Abcam (# ab4674) | 1/200 |
| Rat anti-IBA1 | Wako Chemicals (# 019-19741) | 1/200 |
| Mouse monoclonal IgG1 Anti-6E10 | BioLegend (# 803001) | 1/200 |
| **Immunoblotting** | | |
| Mouse monoclonal IgG1 anti-FLAG® M2 | Sigma-Aldrich (# F1804) | 1/5000 |
| Rabbit anti-PSAP | ProteinTech (# 10801-1-AP) | 1/1000 |
| Rabbit monoclonal anti-BDNF | Abcam (# ab108319) | 1/1000 |
| Rabbit anti-NTF4 | ProteinTech (# 12297) | 1/1000 |
| Mouse IgG2b monoclonal anti-SNAP25 | ProteinTech (# 60159) | 1/1000 |
| Rabbit anti-synaptophysin 1 | ProteinTech (# 17785) | 1/2000 |
| Rabbit anti-synaptotagmin 1 | ProteinTech (# 14511) | 1/500 |
| Rabbit anti-synapsin I | ProteinTech (# 20258) | 1/1000 |

| | | |
|---|---|---|
| Rabbit anti-PSD95 | ProteinTech (# 20665) | 1/1000 |
| Rabbit monoclonal anti-SAP97 | Abcam (# ab300481) | 1/1000 |
| Mouse monoclonal IgG1 anti-SAP102 | BioLegend (# 832002) | 1/1000 |
| Mouse monoclonal IgG1 anti-GluA1 | Antibodies Incorporated (# 75-327) | 1/1000 |
| Mouse monoclonal IgG anti-ACTB | Abcam (# ab8226) | 1/10000 |
| Mouse monoclonal IgG2a anti-saposin C | Santa Cruz (# sc-374119) | 1/1000 |
| Chicken anti-β tubulin 3 | MyBioSource (#MBS 835547) | 1/1000 |

#### Supplemental Figure 1

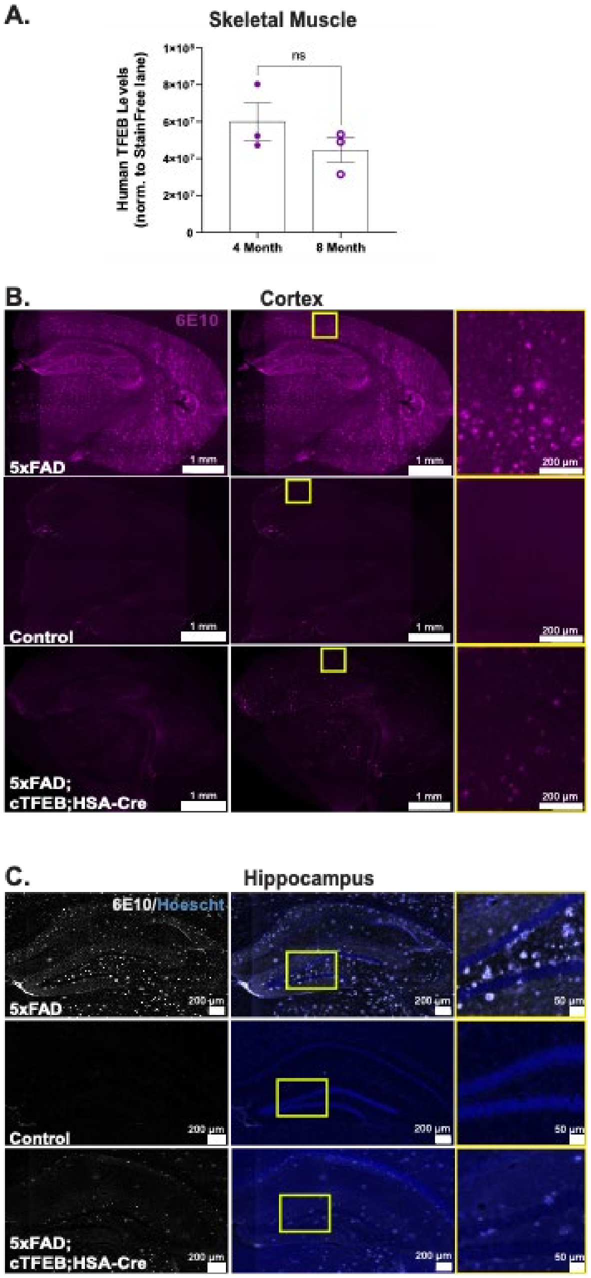


**FIGURE S1.** Densitometry quantification of 3x-FLAG-TFEB (**Figure 1B**) normalized by Total StainFree protein lane densitometry. Closed and open circles represent 4- and 8-month-old 5xFAD;cTFEB;HSACre female transgenic mice. **(B)** Representative merged images of 8-month-old cortex stained for Aβ plaques (6E10, magenta). **(C)** Representative merged images of 8-month-old hippocampus stained for Aβ plaques (6E10, white). Scale bars as shown. Statistical comparison was performed using t-test, n.s. non-significant. Data is represented as mean ± SEM.

### Supplemental Figure 2

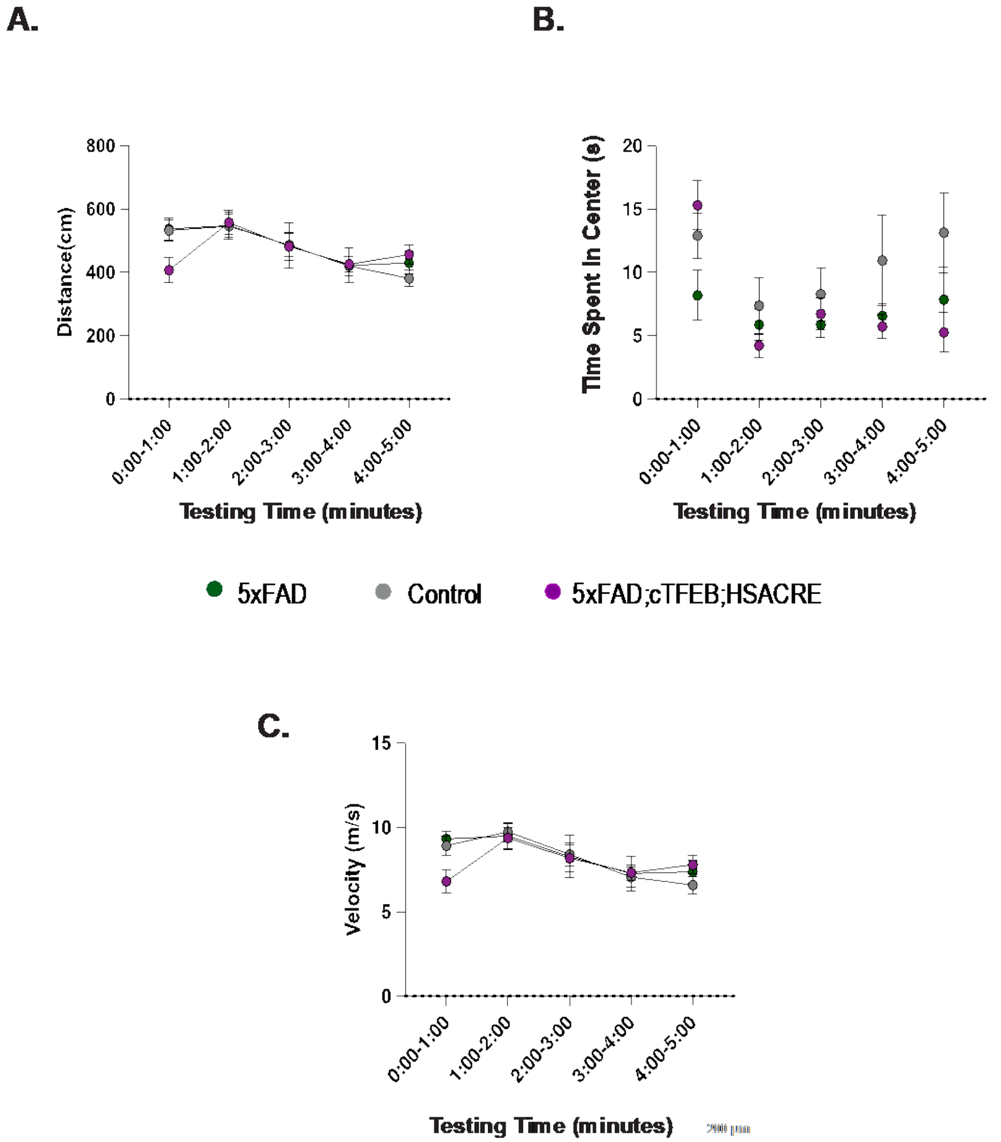


**FIGURE S2. Skeletal muscle TFEB did not alter overall locomotor activity of 5xFAD 8-month-old transgenic female mice. (A)** Open field test evaluating distance traveled, velocity and time in center as proxies for activity levels (n = 8–12/group). Statistical comparison was performed using two-way ANOVA and post hoc multiple comparisons, lack of annotation indicates comparisons were not significant. Data is represented as mean ± SEM.

#### Supplemental Figure 3

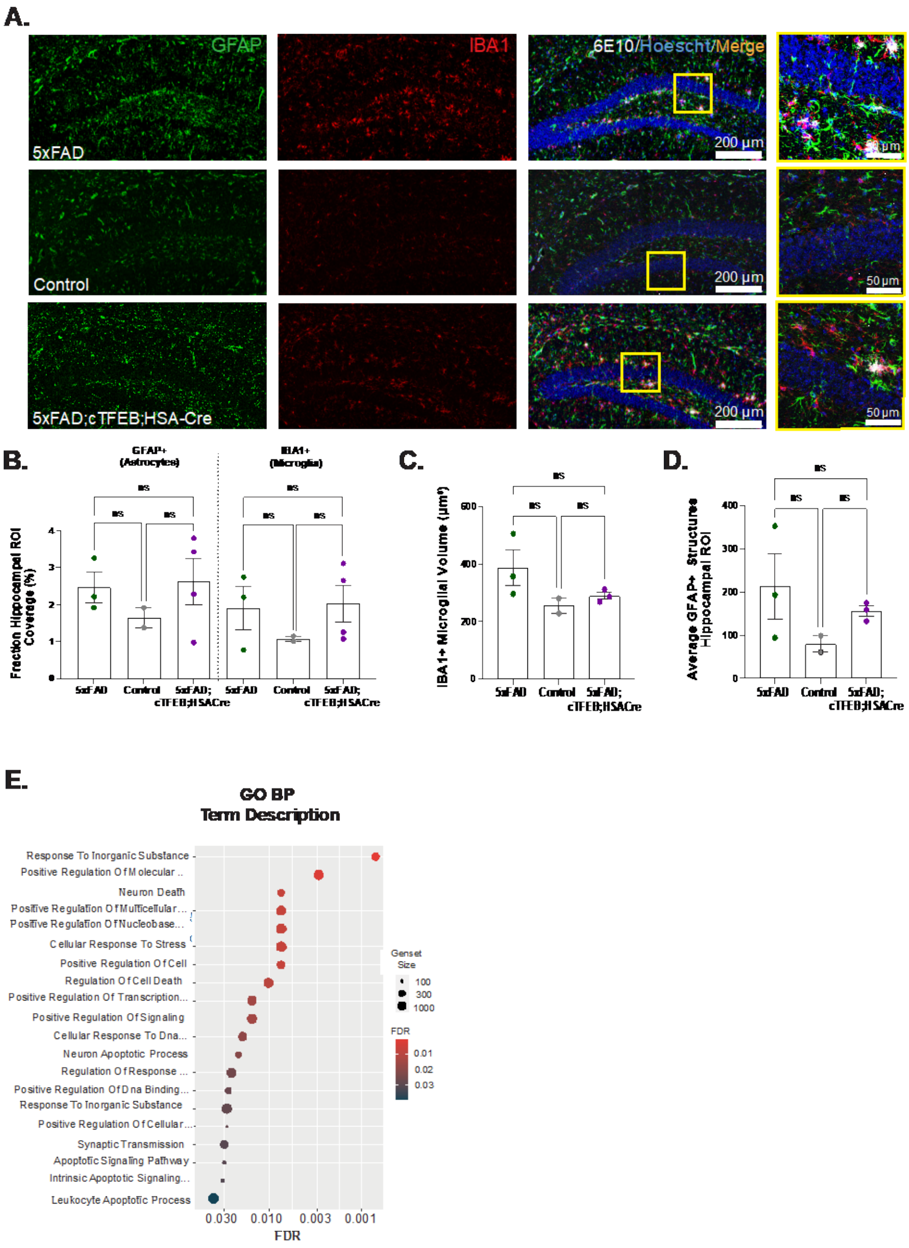


**FIGURE S3. Skeletal muscle TFEB overexpression did not alter neuroinflammation in 4-month-old 5xFAD transgenic female mice. (A)** Representative merged images of the dentate gyrus stained for astrocytes (GFAP, green), microglia (IBA1, red), 6E10 (white) and Hoechst (blue). Scale bars as shown. **(B)** Quantification astrocyte and microglia load. **(C)** Volume of IBA1+ microglia. **(D)** Number of GFAP+ astrocytes. **(E)** Dot plots of the gene ontology (GO) enrichment analysis for transcripts involved in biological processes according to a neuroinflammation Nanostring panel comparing 5xFAD;cTFEB;HSACre vs. 5xFAD 8-month-old female transgenic mice (see **Figure 2E**). Statistical comparison was performed using one-way ANOVA and post hoc multiple comparisons, n.s. non-significant.

#### Supplemental Figure 4

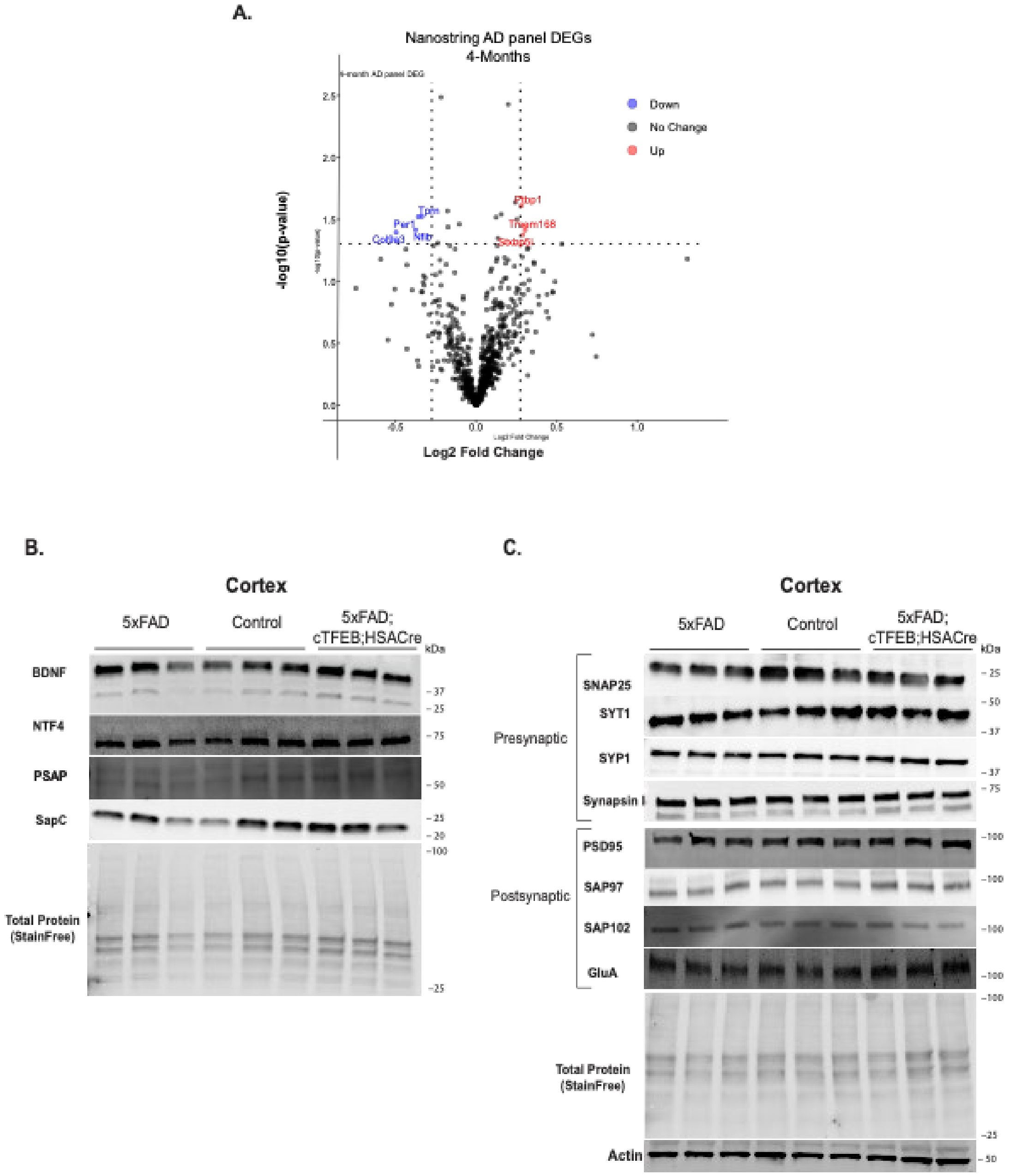


**FIGURE S4. Skeletal muscle-TFEB overexpression did not alter neurotrophic signaling or synaptic integrity in brains of 5xFAD 4-month-old transgenic female mice. (A)** Volcano plot of differentially expressed hippocampal transcripts quantified using a Nanostring AD panel for comparing 5xFAD;cTFEB;HSACre vs. 5xFAD 4-month-old transgenic female mice. **(B-C)** Immunoblot for several neurotrophic **(A)**, pre- and post-synaptic **(B)** markers using 4-month-old mice cortical protein lysates. Total StainFree protein lane or actin β band was used a loading control.

#### Supplemental Figure 5

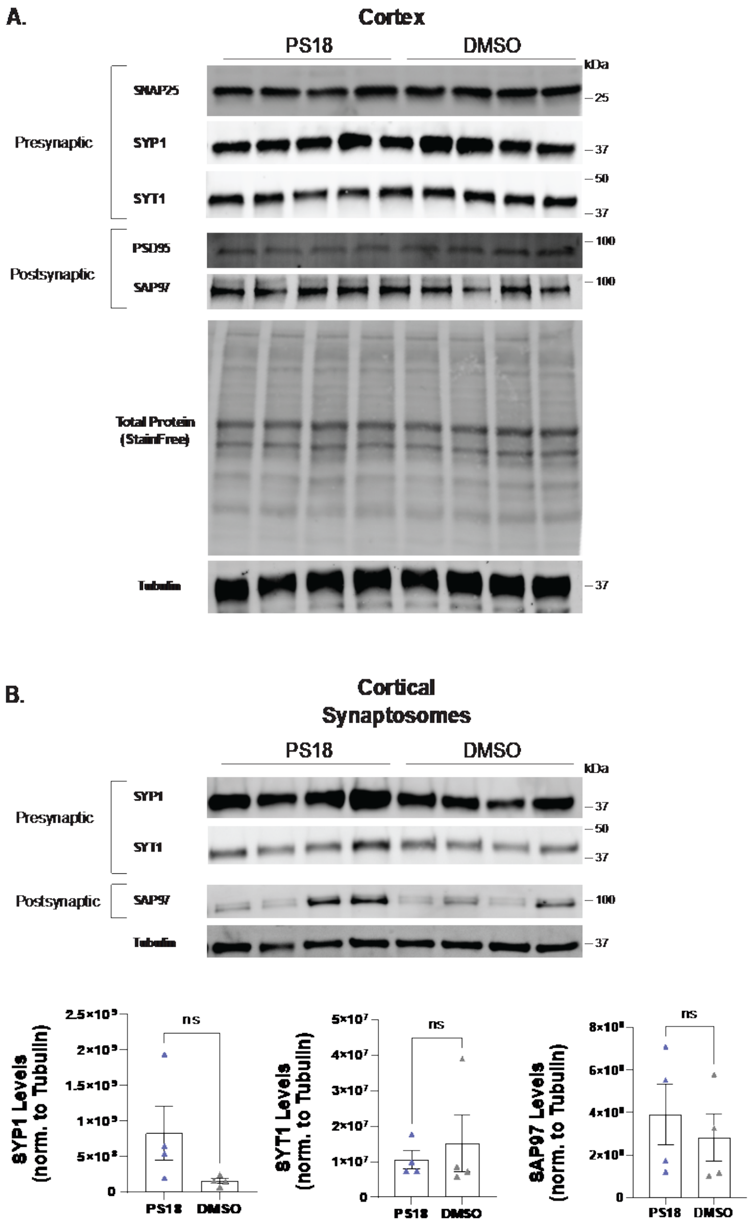


**FIGURE S5. No changes in synaptic markers in total cortical lysates and trending alterations in cortical synaptosomes preps from PS18-injected animals. (A)** Immunoblots of total cortical protein lysates for pre- and post-synaptic markers in subcutaneously PS18- or DMSO-injected 3-month-old C57B6 /L male mice. Total StainFree protein lane or by β tubulin 3 band was used a loading control. **(B)** Immunoblots of cortical synaptosome-enriched protein fractions for pre- and post-synaptic markers in subcutaneously PS18- or DMSO-injected 3-month-old C57B6 /L male mice. β tubulin 3 band was used a loading control. Densitometry quantifications are shown below. Statistical comparison of groups was performed using t-test, n.s. non-significant.
